# Acute resistance exercise induces mitophagy and mitophagomes on subsarcolemmal clefts in human skeletal muscle: Focus on BNIP3L/NIX40 as a mitophagy flux marker

**DOI:** 10.1101/2023.05.22.541782

**Authors:** F. Díaz-Castro, M. Tuñón-Suárez, N. Quezada, P. Rivera, J. Cancino, H. Zbinden-Foncea, E. Morselli, M. Castro-Sepúlveda

## Abstract

**Objective:** Mitochondrial dynamics and quality control in skeletal muscle are central to healthy metabolism. Resistance exercise is a recognized tool for improving skeletal muscle function; however, its effect on mitochondria is still not fully understood. We investigated the impact of resistance exercise on mitochondrial morphology and mitophagy in human skeletal muscle.

**Methods:** Eight healthy men performed resistance exercise on one leg, and muscle biopsies were subsequently obtained from the resting leg (Rest) and the exercised leg (Ex) to measure protein abundance and mitochondrial morphology. Additionally, muscle biopsies were obtained from twelve healthy men, and the abundance of BNIP3L protein was correlated with muscle cell and whole body health markers.

**Results:** Ex increased p-Drp1 and decreased MFN2, Parkin, and BNIP3L protein levels. Electron microscopy indicated an increase in mitochondrial circularity, cristae abnormality, and mitophagosome structures in Ex, with a marked increase in subsarcolemmal mitophagosomes. We also identified mitophagosomes outside the muscle. Experiments in human myotubes showed a severe decrease in BNIP3L protein in response to CCCP-induced mitochondrial damage with and without bafilomycin. A positive correlation was found between BNIP3L and exercise RQ, HOMA index, while a negative correlation was found with mitophagosomes abundance and VO_2_max.

**Conclusion:** Our study describes the effect of resistance exercise on mitochondrial dynamics and mitophagy in skeletal muscle, demonstrating induction of mitochondrial fission and mitophagy in the exercised leg. Moreover, we propose BNIP3L as a potential regulator and marker of mitophagy flux.

## 1. Introduction

Mitochondria adapt their morphology and function in response to intracellular and extracellular stimuli [1]. Their morphology depends on the dynamic processes of mitochondrial fusion and fission. Mitochondrial fusion is regulated by MFN1/2 and OPA1, connecting outer and inner mitochondrial membranes, respectively [2]. Mitochondrial fission is regulated by Drp1, which generates a constricted ring around the mitochondria driving its fission [3]. Through fission dysfunctional mitochondria, presenting low membrane potential or irreversible mtDNA damage, are separated to be degraded by autophagy (mitophagy) [4,5]. Mitophagy, serves as a vital quality control mechanism to preserve healthy mitochondria to maintain cellular energy homeostasis [6]. PINK1/Parkin proteins have been described as central mitophagy regulators, however, mitophagy receptors, such as BNIP3L/NIX40 and FUNDC1 have been more recently identified as additional mitophagy regulators [7,8].

The skeletal muscle (SkM), which accounts for around 40% of body mass, is highly metabolically active in the body human and plays a crucial role in insulin-stimulated glucose disposal [9]. Mitochondria play an essential role in SkM metabolism, and an unbalance in mitochondrial dynamics has been related to lipotoxicity and insulin resistance [10,11] as well as sarcopenia and dystrophy in humans [12,13], placing mitochondrial dynamics as a critical process in SkM homeostasis. Mitophagy decrease causes the accumulation of damaged mitochondria and the deterioration of mice SkM [14]. Therefore, enhancing mitophagy in SkM could improve SkM quality and metabolic health in humans.

Exercise improves muscle metabolism and reduces metabolic disorders [15]. In old men regular aerobic training increases mitophagy in human SkM, as suggested by the increase in mitophagy proteins together with an increase in mitochondrial metabolic function [16]. Resistance exercise is a recognized tool to improve SkM mass and contractile function [17], however, whether this kind of exercise influences human SkM mitochondrial dynamic and/or mitophagy has yet to be established [18]. Resistance training increases mitochondrial oxidative capacity in human SkM, however, the effect on mitochondrial density is inconsistent [19–21], suggesting that resistance training can increase mitochondrial function without affecting mitochondrial density. In rat, resistance exercise mimetic induces mitochondrial fission in SkM, as indicated by the increase in Drp1-ser616 phosphorylation [22], an initial step in mitophagy. We hypothesize that resistance exercise increases fission and mitophagy, promoting the degradation of damaged mitochondria in human SkM. In support of this hypothesis, a recent study shows that a single bout of resistance exercise induces mitochondrial cristae damage in SkM (which indicates the beginning of mitophagy), and chronically increases mitochondrial cristae density [23], a marker of increased metabolic capacity [10,24]. Therefore, the aim of this study is to elucidate the effects of resistance exercise on mitochondrial fission and mitophagy.

## 2. Methods

### 2.1. Subjects and protocol study 1

Eight nondiabetic, nonsmoking men, without a history of cardiovascular, respiratory, or thyroid disease, were included. <u>First appointment</u>: Subjects after an 8-h fast, without consuming caffeine or alcohol for 48-h, were evaluated to measure one repetition max (1RM), weight, and height. <u>Second appointment</u>: Subjects after an 8-h fast, without consuming caffeine or alcohol for 48-h, performed the resistance exercise protocol in the dominant leg. At the end of the exercise a biopsy from the Vastus Lateralis muscle of the resting leg (Rest) and exercised leg (Ex) was taken. The biopsy was split into two pieces: (1) Immediately frozen in liquid nitrogen and stored at -80°C for western blotting and histology, and (2) fixed in 2.5% glutaraldehyde for transmission electron microscopy (TEM) analyses. All subjects signed a written informed consent approved by the Institutional Review Board of the Universidad Finis Terrae and adhered to the Declaration of Helsinki.

### 2.2. Evaluation of one repetition max (1RM) and training session

Evaluation of 1RM was performed in a leg press machine. A warm-up of 8-10 repetitions with 50% of the 1RM estimate was performed. Next, 5 repetitions with 70% estimate 1RM, 3 repetitions with 80% estimate 1RM, and 1 repetition with 90% estimate 1RM. 2-3 kilograms were increased after each repetition until reaching the repetition with the maximum possible weight. Subjects rested between 3-5 minutes between stages. 1RM was registered as the final weight lifted completely. After one week of the 1RM evaluation, the voluntaries arrived at the laboratory at 7:00 AM following an 8-h fast, without previous consumption of caffeine/alcohol, and after warming up in the leg press machine (10 repetitions with 22kg), voluntaries performed 10 series of 10 repetitions at 70% of 1RM with 2 minutes rest between series. Biopsies were taken within 1-hour time, at the end of the exercise.

### 2.3. Western blot

Muscle biopsies were homogenized using an electric homogenizer in a buffer containing: 20 mM of Tris-HCl (pH 7.5), 1% of Triton X-100, 2 mM of EDTA, 20 mM of NaF, 1 mM of Na2P2O7, 10% of glycerol, 150 mM of NaCl, 10 mM of Na3VO4, 1 mM of PMSF, and a protease inhibitor cocktail (complete mini; Roche Applied Science). Proteins were separated by SDS-PAGE and transferred to PVDF membranes, as previously described [25]. The following antibodies (dilutions) were used: Mfn2 (1:1000, ab56889, Abcam), Opa1 (1:1000, 612606, BD Biosciences), Total OXPHOS cocktail (1:1000, ab110413, Abcam), LC3 (1:1000, 4108, Cell Signaling Technology), p-mTOR (1:1000, 2971, Cell Signaling Technology), p-AMPK (1:1000, 2531, Cell Signaling Technology), p62 (1:2000, H00008878-M01, Abnova), p-Drp1ser616 (1:1,000, SC-101270, Santa Cruz Biotechnology), PINK1 (1:1000, NB.BC100-494, Novus Biological), Parkin (1:1000, SC-32282, Santa Cruz Biotechnology), BNIP3L (1:1000, 12396, Cell Signaling Technology), FUNDC1 (1:1000, ab224722, Abcam). GAPDH (1:10000, 2118, Cell Signaling Technology) was used as loading control. Protein bands were visualized on a film (WESTAR SUPERNOVA detection kit, Cyanagen, Bologna, Italy), scanned, and quantified by densitometry using Image Lab Software Version 6.0 (Bio-Rad).

### 2.4. Histology

A piece (∼50 mg) of the muscle biopsy was immediately frozen and stored at −80°C. Cryosections (10 μm) of muscles were stained with succinate dehydrogenase (SDH) following standard protocols. SDH was quantified in ∼70 fibers/volunteer using Fiji/ImageJ software.

### 2.5. Transmission Electron Microscopy (TEM) sample preparation

Biopsies were fixed in a glutaraldehyde solution (2.5%, room temperature, 2 hours). Fixed muscle was dissected into bundles of fibers, washed four times with 0.1 M sodium cacodylate buffer, and stained with 2% osmium tetroxide in 0.1 M sodium cacodylate buffer for 2 hours. Samples were washed with water, stained with 1% uranyl acetate for 2 hours, dehydrated on acetone dilution series, and embedded in Epon resin. Finally, 80-nm sections were cut, mounted on grids, and examined using a TEM (Talos F200C G2; Microscopy Facility, Pontificia Universidad Católica de Chile).

### 2.6. TEM image analyses

For mitochondrial morphology, a mask was drawn over the outer membrane of the mitochondria at ×15000 magnification, as described [25,26]. The mitophagosomes were identified as previously described [27]. TEM images were analyzed on longitudinal sections using Fiji/ImageJ software.

### 2.7. Immunofluorescence

Muscle sections of 30 μm thickness were made using a cryostat (Leica Biosystems, IL, USA) at −20 °C. Fixed muscle sections were blocked with 3% bovine serum albumin (BSA, Winkler, Chile) in phosphate buffered saline (PBS) and permeabilized with 0.25% Triton X-100 for 1 h at room temperature. Then, brain sections were incubated overnight at 4 °C with the primary antibody against TOM20 (sc-136211, Santa Cruz) and LC3 (4108, Cell Signaling Technology) prepared in 0.05% Triton X-100 and 3% BSA in PBS, followed by conjugation with its secondary antibody (Alexa Fluor, Life Technologies, Carlsbad, CA, USA), for 1 h at room temperature. Confocal fluorescence microscopy assessments were performed using an LSM 880 Zeiss inverted confocal microscope with Airyscan detection (Unidad de Microscopía Avanzada UC (UMA UC). Contrast and/or brightness adjustment and cropping of final images were done using ImageJ, with identical settings applied to all images from the same experiment.

### 2.8. Human myotubes treatment

A pool (n=3) of myoblasts from healthy humans was differentiated as described previously [28] to obtain myotubes. Myotubes were treated with palmitate (300uM), CCCP (10uM) and bafilomycin (100nM) for 24h. Then cells were lysated with T-per buffer (78510, Thermo Fisher Scientific) supplemented with protease inhibitor cocktail (complete mini; Roche Applied Science). Western blot was performed as described in section 2.3.

### 2.9. Subjects and protocol study 2

We analyzed a new subset of data from a previous study of our group [25]. VO_2_max, HOMA, cross-sectional area (CSA), force, and resting/exercise respiratory quotient has been previously published [25] but herein we reanalyzed those data differently.

### 2.10. Statistics

Data are presented as means [standard error of the mean]. T-test was performed for two groups analysis and P<0.05 was considered statistically significant. Correlations between study variables were assessed with the Pearson test, assuming a normal distribution of the data, and P<0.05 was considered statistically significant. Prism 9 Analyses were performed using Prism 9 (GraphPad Software, La Jolla, CA)

## 3. Results

### 3.1. One resistance exercise bout increases mitochondrial fission proteins and decreases mitophagy proteins in human skeletal muscle

After a single bout of resistance exercise, we found in the exercised leg, hereafter Ex, an increase in the phosphorylation of p-Drp1ser616, a trend to decrease Mfn2, and no changes in the mitochondrial fusion OPA1 in comparison to the resting leg, hereafter Rest (Figure 1B). Among Oxidative Phosphorylation (OXPHOS) proteins, only complex IV decreases in Ex when compared to Rest (Fig1E). SDH activity (complex II) does not change in Ex (Fig1G). The autophagy markers LC3 and p62 (Fig1C), as well as AMPK phosphorylation in threonine 172 do not change (Fig1C). Despite this mTOR phosphorylation in serine 2448, a master regulator of the autophagy pathway increased in Ex. PINK1 and FUNDC1 levels are not affected by Ex, however, Parkin and BNIP3L levels are decreased (Fig1D).

**Figure 1.**
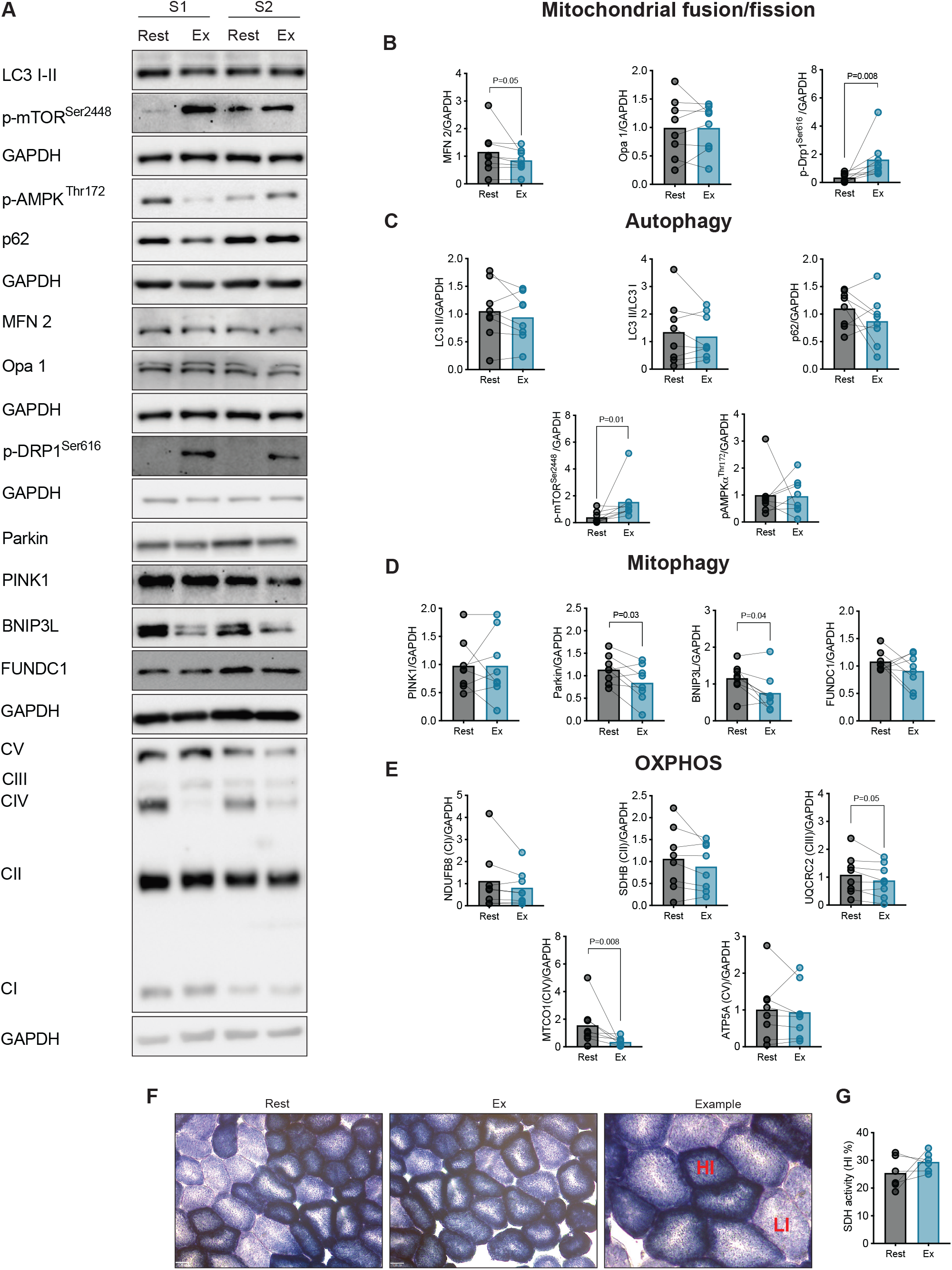
Resistance exercise increases p-Drp1^Ser616^ and decreases mitophagy proteins in human skeletal muscle. (A) Representative image of the measured proteins of two participants in resting (Rest) and exercised (Ex) legs biopsies. (B) Densitometry of western blot proteins related to mitochondrial fusion and fission. (C) Densitometry of western blot proteins related to autophagy. (D) Densitometry of western blot proteins related to mitophagy. (E) Densitometry of western blot proteins related to oxidative phosphorylation (OXPHOS). (F) Representative image of succinate dehydrogenase (SDH) staining in Rest and Ex legs. HI: High intensity; LI: Low intensity. (G) Quantification of SDH staining. Statistical significance was tested by a paired t-test. P <0.05 was considered statistically significant, while a p-value between 0.05 and 0.09 was considered a trend. The dots in the bar graphs represent different participants, and the data are shown as means.

### 3.2. Resistance exercise increases mitochondrial abnormal cristae and mitophagosomes in human skeletal muscle

We evaluated mitochondrial morphology through transmission electron microscopy in intermyofibrillar (IMF) and subsarcolemmal (SSM) mitochondria. Mitochondrial size did not change (Figure 2C, D), however mitochondrial elongation decreased on Ex in both mitochondrial subpopulations in comparison to Rest. Abnormal mitochondrial cristae are increased in both IMF and SSM, accompanied by an increase in mitophagosomes-like structure in Ex leg (Figure 2C, D). Also, we observed more mitophagosomes-like structures on the cleft in the subsarcolemmal membrane on Ex (Figure 2E) in comparison to Rest. Interestingly, we identified mitophagosome-like structures outside the muscle in Ex group (Figure 2E). In an immunofluorescence performed on 1 subject, we observed an increase in LC3 mark in Ex condition (Figure 2F)

**Figure 2.**
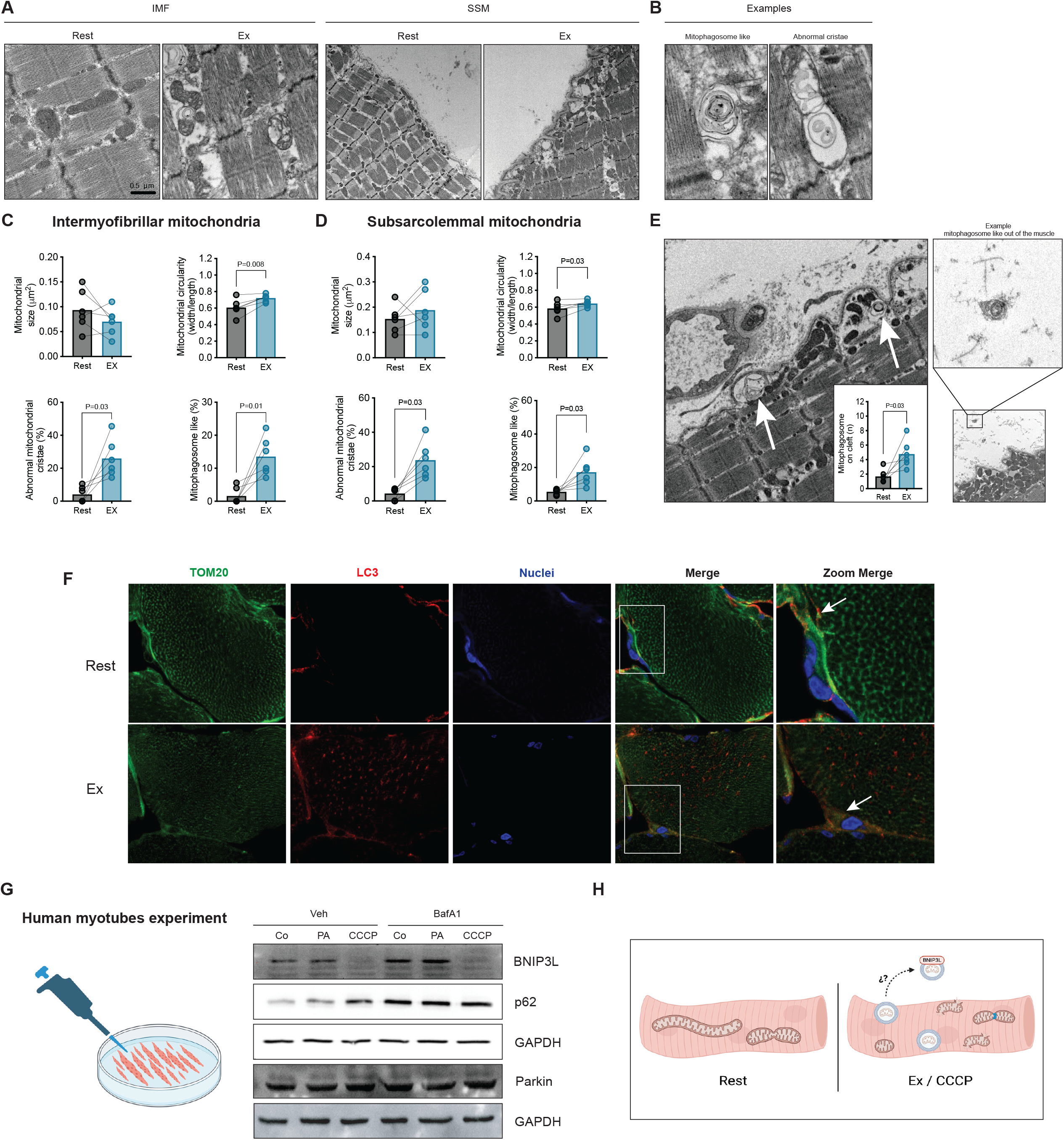
Resistance exercise induces a fragmented mitochondrial phenotype and increases mitochondria damage and mitophagosomes. (A) Transmission electron microscopy representative image of intermyofibrillar (IMF) and subsarcolemmal (SSM) mitochondria. (B) Representative image of mitophagosomes and abnormal cristae. (C) Mitochondrial morphology and mitophagosome quantification of IMF. (D) Mitochondrial morphology and mitophagosome quantification of SSM. (E) Representative image and quantification of mitophagosome on cleft. Example of mitophagosome-like out. (F) Representative immunofluorescence of TOM2 (green), LC3 (red), and nuclei (blue) of 1 subject. White area in merge is represented in zoom merge. White arrow show LC3 and TOM20 mark in membrane. (G) Representative western blot image of BNIP3L, p62, parkin, and GAPDH of human myotubes stimulated with palmitic acid (PA), Carbonyl cyanide 3-chlorophenylhydrazone (CCCP) or bafilomycin (BafA1). (H) Graphical hypothesis proposal of mitochondria expelled from the skeletal muscle mediated by BNIP3L. P <0.05 was considered statistically significant, while a p-value between 0.05 and 0.09 was considered a trend. The dots in the bar graphs represent different participants, and the data are shown as means.

### 3.3. BNIP3L decreases severely in response to mitochondrial damage

To test if BNIP3L and Parkin respond to mitochondrial damage, human myotubes from healthy men were stimulated with palmitate, CCCP, and BaFA1, this last to inhibit lysosomal degradation. BNIP3L decreased dramatically with CCCP but not after palmitate exposure. BNIP3L also decreased following CCCP treatment in the presence of BaFA1. The autophagy marker p62 increased with CCCP and accumulated in the presence of BaFA1. The different treatments did not affect Parkin levels (Figure 2G).

### 3.4. BNIP3L is associated with exercise RQ and insulin resistance

Since BNIP3L was the only protein that decreased in response to mitochondrial damage (resistance exercise and CCCP), we hypothesize that BNIP3L could be a marker of mitophagy flux. Our data show an inverse correlation between BNIP3L and mitophagosome percentage and cristae density (Figure 3C, D). Additionally, BNIP3L abundance in human SkM is positively associated with lipid droplet density (trend) and exercise RQ (Figure 3E, F). Also, an inverse correlation is observed between BNIP3L with VO_2_max and HOMA (Figure 3F). No correlation was found between BNIP3L and CSA, force, and resting respiratory quotient (Data not shown).

**Figure 3.**
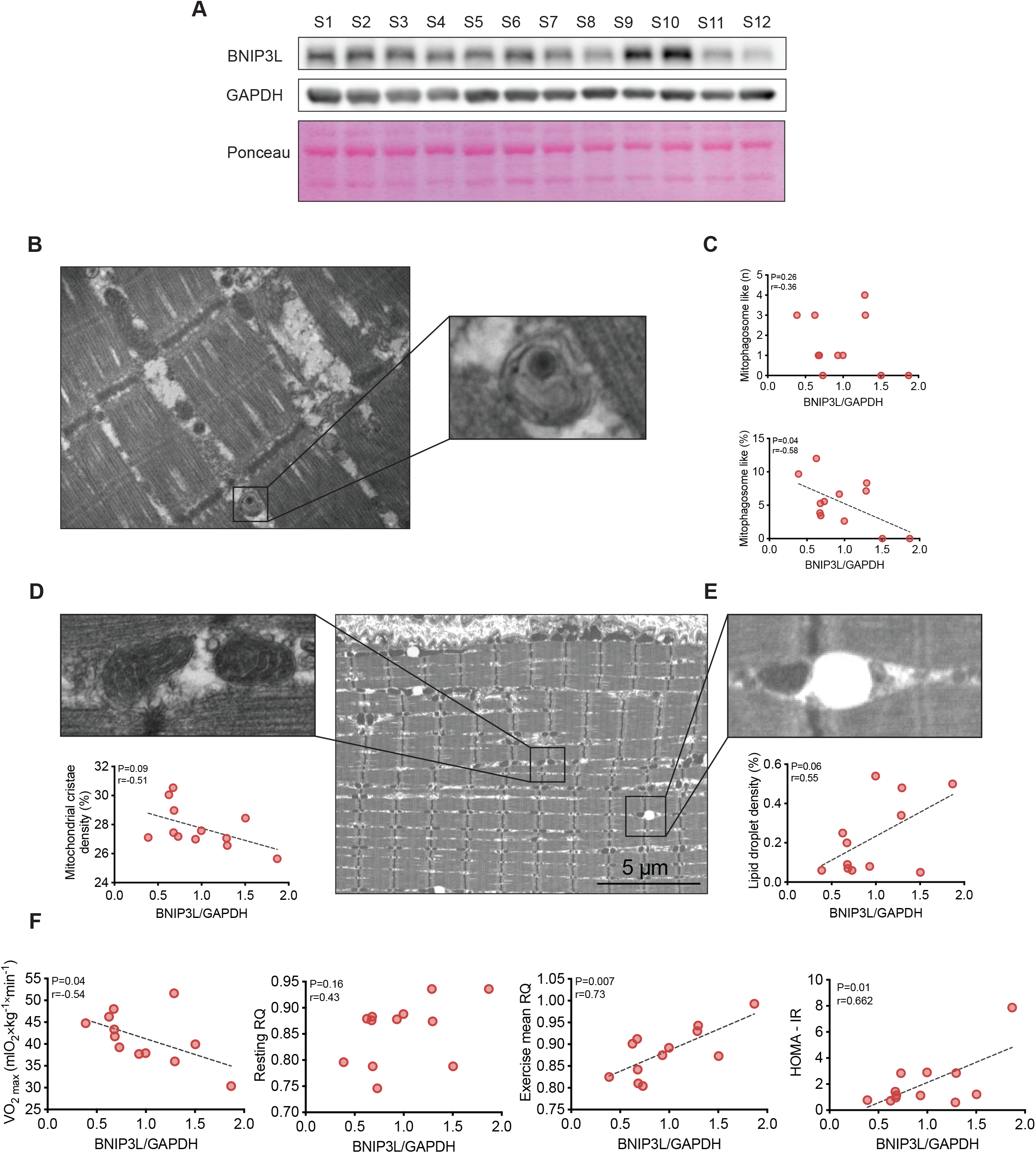
BNIP3L in human skeletal muscle correlate with mitophagosome abundance, metabolic capacity, fatty acid oxidation during exercise, and insulin resistance. (A) Representative image of BNIP3L western blot of twelve participants. (B) Representative image of mitophagosomes. (C) Correlation of mitophagosomes number with percentage with BNIP3L. (D) Representative image of mitochondria cristae and its correlation with BNIP3L. (E) Representative image of lipid droplet and its correlation with BNIP3L. (F) Correlation between BNIP3L with VO_2_max, resting respiratory quotient (RQ), exercise mean respiratory quotient, and HOMA. Correlations were made with Pearson. P <0.05 was considered statistically significant, while a p-value between 0.05 and 0.09 was considered a trend. The dots in the bar graphs represent different participants.

## 4. Discussion

Here we investigated the effects of a single resistance exercise bout on mitochondrial dynamics in human SkM using a single-leg model, hypothesizing that resistance exercise induces mitochondrial fission and mitophagy. Our results show that a single bout of resistance exercise increases the mitochondrial fission protein p-Drp1ser616 and decreases the mitophagy proteins Parkin and BNIP3L. Ex increase mitochondrial fragmentation, abnormal mitochondrial cristae, and mitophagosomes-like structures. Moreover, we observed mitophagosome-like structures outside of the SkM in Ex. In an in vitro model we find that BNIP3L protein decreases severely in response to mitochondrial damage, independent of lysosomal degradation, suggesting that mitochondria could be secreted out of the SkM. To gain insights into the relevance of the BNIP3L protein in human SkM, we correlated BNIP3L protein abundance from biopsies with muscle cell markers and general whole-body health markers, identifying a positive correlation between BNIP3L with exercise RQ, HOMA, while a negative correlation with mitophagosome and VO_2_max.

With respect to autophagy, we did not observe differences in LC3 nor p62 in response to resistance exercise however previous reports show a decrease in LC3-II protein levels 1-3 hours after resistance exercise with no changes in p62 protein levels [29,30]. Differences in the exercise protocol could explain the observed difference (Two legs exercise with pre-post biopsy vs our single-leg protocol), suggesting that plasmatic factors or metabolic changes induced by resistance exercise could be more relevant than a mechanical stimulus on autophagy induction. Mitochondrial fission is necessary for mitophagy, and our results suggest that mitochondrial damage (abnormal cristae), fission phenotype (p-Drp1ser616 and mitochondrial circularity), and an increase in mitophagy (increases in mitophagosome-like structures and decreases in mitophagy proteins) are induced by resistance exercise. Botella et al., in a recent study, showed a low aspect ratio, more circularity, and an increase in mitochondrial damage after a resistance exercise bout in strength athletes [23]. However, in another study, no differences were found after 24 hours of resistance exercise in mitochondrial fission proteins [31]. We speculated that resistance exercise forces mitophagy, removing damaged mitochondria in SkM, and consistently with our hypothesis, we saw an increase in intermyofibrillar on cleft and outside mitophagosomes, with a decrease in BNIP3L and Parkin proteins. The possibility that mitochondria can be expelled from the cell has been explored in other models [32–34], however, this is the first time it has been observed in human SkM after exercise. Through in vitro experiments, we could show that BNIP3L is the only mitophagy marker that decreases in response to mitochondrial damage independent of lysosomal degradation, hence we propose that BNIP3L could play a role in mitochondrial expulsion from the human SkM after mitochondrial damage.

BNIP3L, also known as NIX40, is a mitophagy receptor of the BCL-2 family involved in the regulation of apoptosis, necrosis, autophagy, and mitophagy [35,36]. Its role in the maintenance of mitochondrial health in SkM has been well reported in rodent and in vitro models [7,37], whether this is maintained also in human SkM is still unknown. As BNIP3L responds to mitochondrial damage, both in resistance exercise and in CCCP treatment, we correlated the abundance of BNIP3L in human SkM with health parameters finding a direct correlation with exercise RQ and HOMA, indicating that lower BNIP3L protein expression in SkM is associated with higher fatty acid oxidation during exercise and insulin sensibility, respectively. One possible explanation for these results is that BNIP3L could be used as a marker of mitophagy flux. The direct correlation between BNIP3L abundance and the percentage of mitophagosome-like structures suggests that BNIP3L accumulation coincides with the number of mitophagosomes. However, in conditions where mitophagy is increased, such as during resistance exercise, BNIP3L levels decrease. We speculate that BNIP3L accumulation may indicate the number of damaged mitochondria. To confirm this hypothesis, additional experiments are needed. Moreover, limitations of our study, such as ensuring the veracity of mitophagosomes and determining if the mitochondrial output in SkM is for a BNIP3L mechanism or membrane damage, must be considered in future investigations.

This is the first study that investigates the effect of resistance exercise in mitophagy on human SKM. In conclusion, our results show that resistance exercise induces mitochondrial fission and mitophagy in human SkM, in addition, our findings suggest that BNIP3L protein levels correlate with metabolic health and could be a mitophagy flux marker in human SkM. We hypothesize a new possible mechanism of mitochondrial clearance, independent of lysosomal degradation, mediated by BNIP3L in SkM, however, more experiments as mitochondria tracking to confirm their expulsion from the SkM or the BNIP3L deletion in a resistance exercise protocol, are needed to fully understand the role of BNIP3L in mitochondrial response and adaptation to resistance exercise.

## 5. Author contributions

MC-S conceived the study. MC-S, FD-C, and MT designed the study. MC-S, FD-C, MT, NQ, PR, and JC executed experiments and processed and analyzed data. FD-C, MC-S, and EM drafted the manuscript; FD-C, MC-S, EM, and HZ revised the manuscript and contributed significantly with intellectual content. All authors approved the submitted version of the manuscript.

## 6. Data availability

Data will be made available on request.

## 7. Acknowledgements and funding

Thanks to all participants. This work was supported by the Unidad de Microscopía Avanzada UC (UMA UC). This study was funded by grants from Universidad Finis Terrae (Grant no. CAI 2021 and CICI 2022), Agencia Nacional de Investigación y Desarrollo (ANID), Chile: Doctorado Nacional 21210611 (FD-C), 21211189 (PR); FONDECYT 1200499 (EM), 11230548 (MC-S).

